# Protective microbiomes can limit the evolution of host pathogen defense

**DOI:** 10.1101/665265

**Authors:** C. Jessica E. Metcalf, Britt Koskella

## Abstract

The evolution of host immunity occurs in the context of the microbiome, but little theory exists to predict how resistance against pathogens might be influenced by the need to tolerate and regulate commensal microbiota. We present a general model to explore the optimal investment in host immunity under conditions in which the host can, versus cannot easily distinguish among commensal versus pathogenic bacteria; and when commensal microbiota can, versus cannot protect the host against the impacts of pathogen infection. We find that a loss of immune vigilance associated with innate immunity over evolutionary time can occur due to the challenge of discriminating between pathogenic and other microbe species. Further, we find the greater the protective effect of microbiome species, acting either directly or via competition with a pathogen, or the higher the costs of immunity, the more likely the loss of immune vigilance is. Conversely, this effect can be reversed when pathogens increase host mortality. Generally, the magnitude of costs of immunity required to allow evolution of decreased immune vigilance are predicted to be lowest when microbiome and pathogen species most resemble each other (in terms of host recognition), and when immune effects on the pathogen are weak. Our model framework makes explicit the core trade-offs likely to shape the evolution of immunity in the context of microbiome / pathogen discrimination. We discuss how this informs interpretation of patterns and process in natural systems, including vulnerability to pathogen emergence.

**Impact Summary:** Evidence for impacts of the microbiome on host health is accumulating. Despite this, little theory has been developed to delineate the evolutionary trajectories that might lead to observed host-microbiome associations. One particularly important theoretical gap is evaluating how the presence and effects of microbiome species modify selection pressure on immune system function. We develop a simple model of host fitness given both immune discrimination and microbiome and pathogen effects on survival, in the context of an interaction between the microbiome and pathogen species. We use this framework to predict when and to what degree the presence of microbiome species might lead to loss of immune vigilance. Positive microbiome effects can drive loss of immune vigilance, whether the microbiome acts directly on pathogen growth or indirectly by reducing the impacts of pathogens; and high costs of immunity will amplify this effect. Our results provide a first set of predictions regarding how immunity should evolve given the challenge of discriminating pathogen and microbiome species, and reveals the ways in which this might leave hosts vulnerable to novel pathogens.

## Introduction

Bacterial species making up the microbiome are increasingly recognized to play an important role in host health, including via microbiome-mediated protection against pathogens (Friesen et al. 2011). An important implication is that the evolution of host immunity must have occurred in the context of balancing tolerance of commensals and resisting pathogens (Littman and Pamer 2011). As a result, underlying ecological features of both hosts and bacteria will define how selection shapes host immunity across host generations. For example, microbiome-mediated protection might evolve very rapidly, due to both shorter microbial generation times and greater standing genetic variation. The resulting protection against pathogens enjoyed by the host might hinder the evolution of host genetic resistance (King and Bonsall 2017). Growing evidence also suggests that the microbiome plays a critical role in shaping the breadth and specificity of host immunity (Thaiss et al. 2016). This will also shape selection pressures on host immunity to resist pathogens. Despite increasing recognition of the importance of the microbiome in host ecology and health, mathematical models of core processes remains rare, and there is little theory to guide expectations (Koskella, Hall, and Metcalf 2017).

To adequately reflect natural systems of interest, mathematical models must capture several interlocking aspects of the ecology and co-evolution underpinning microbiome-mediated protection. Here, we focus our investigation on innate immunity, the first line of defense against pathogens, found across species from plants to vertebrates. We ask how the presence of commensal microbiota might shape selection on how hosts trigger innate immunity. Innate immunity would ideally be launched when molecular signatures of pathogens were detected. Since pathogens will rapidly evolve away from any signature that is detectable by innate immunity, and that is not subject to constraints, typically, the molecular signatures used by innate immunity are highly conserved among bacteria. The bacterial flagellin protein is a classic example (Gómez-Gómez and Boller 2002) - this apparently tightly constrained structure is widely used by hosts as a strong trigger of host innate immunity (Felix et al. 1999). Importantly, the result is that pathogens and commensal microbiota are likely to have overlapping expression of molecules used by the host in detection of pathogens (Levy et al. 2018; Vogel et al. 2016). The decision to trigger an immune response must therefore discriminate between the presence of pathogens and (neutral or even beneficial) microbiota despite these similarities. Hosts are left with a fundamental tension between maintaining a diverse microbiome and defending against pathogens despite these similarities. This tension remains poorly understood.

The problem is further complicated by the ability of many commensal microbiota to play a role in host defense by either excluding pathogen colonization and growth, or reducing the impact of pathogens on host health (Snelders et al. 2018). This might (or might not) echo molecular signatures underlying detection by immunity, for example if phylogenetically related pathogenic and commensal bacteria were more likely to compete for shared resources and were recognized by the immune system through shared mechanisms. In this case, reducing immune vigilance may allow for the proliferation of particular commensal microbiota that directly compete with invading pathogens, reducing the need for hosts to invest in immunity (Hrček et al. 2018; Jaenike 2012). Many lines of evidence suggest that triggering an immune reaction is likely to be costly (Hanssen et al. 2004; Sheldon and Verhulst 1996) and thus walking the line between effective pathogen defenses and regulating microbiome homeostasis should be a ubiquitous but non-trivial challenge.

Here, we ask how host innate immunity might evolve in the presence of a commensal microbiota with the potential to diminish pathogen impact on the host via competition. Since many molecules used by hosts to trigger innate immunity are evolutionarily constrained (e.g., flagellin (Gómez-Gómez and Boller 2002)), we initially assume that evolution in pathogen and commensal microbiota communities is negligible. We capture the impacts of pathogen and commensal microbiota on fitness through a simple mapping from their abundance to host survival (rather than developing a complete dynamical description of the interactions between pathogens, commensal microbiota and host immunity; e.g., as in (Leung and Weitz 2019)). To capture the discrimination problem faced by innate immunity, we assume that pathogens and commensal microbiota can be reflected as being distribution along a single axis, representing a range from a ‘small’ to a ‘large’ signal detectable by the immune system, which might reflect aspects such as the number of CpG repeats (detectable by Toll-like-receptor 9 (Pohar et al. 2015)), immunogenically variable aspects of flagellin (Felix et al. 1999; Trdá et al. 2014), or a combined signal integrating presence of highly conserved microbe-associated molecules with functional characteristics indicative of pathogen’s presence (e.g., via NBS-LRR proteins in plants (DeYoung and Innes 2006) and signatures of ‘stress’ or deviation from homeostasis more broadly (Chovatiya and Medzhitov 2014)). From this, we develop a model framework constructed around core principles from epidemiology to better understand the drivers of the evolution of host defense in the context of pathogen-microbiome interactions. We aim in particular to delineate the conditions under which the presence of commensal microbiota may result in loss of immune vigilance altogether. Our results show that loss of vigilance driven by protective microbiota is possible, and its likelihood increases as similarity between the commensal microbiota and pathogen communities increases, and when immune effectiveness against the pathogen is weak. We discuss how evolution by pathogen and commensal microbiota communities might modulate outcomes for the host, and how these consequences might play out across host life histories.

## Methods and Results

### A discrimination trade-off

The problem faced by the immune system in discriminating between ‘good’ commensals and ‘bad’ pathogens is analogous to a classic challenge from epidemiology. Overlap between two categories along a continuum means that choosing where to draw a line to discriminate between them results in a trade-off between falsely allocating individuals to one or the other category (Figure 1). In this example, the two distributions correspond to the community of commensal microbiota (left) and pathogenic microbiota (right); the height of the two distributions reflects the number of individuals in each category corresponding to each value along the continuum (e.g. shared flagellin characteristics). The immune strategy under selection corresponds to the problem of where to draw the line above which an immune reaction is triggered. If we define the commensal microbiota as the community of microbial species with positive effects, ideally, the line would have all individuals from this community to its left, where no immune response is triggered, and all pathogenic microbiota to its right, where immunity acts to clear microbial species. However, the predicted overlap between these two communities means that this would be impossible. Drawing the line to the far left of the plot corresponds to a strategy of ‘total vigilance,’ where every single microbe along the continuum triggers a reaction (with associated costs of immune response). We refer to this as “100% vigilance”. Drawing the line to the far right of the plot corresponds to a strategy of ‘no vigilance’, where no microbe elicits an immune reaction. If there is overlap between the two distributions, drawing the line towards the middle of the plot (e.g., the intermediate strategy illustrated in Figure 1) inevitably mis-classifies some individuals: some members of the commensal microbiota will be above the line, and some members of the pathogenic microbiota below it. The strategy that maximizes fitness will balance costs of immunity with costs and benefits of pathogens and commensal microbiota, and the effect of their overlap, i.e., the impact of the commensal microbiota on the pathogen relative to outcomes for the host.

**Figure 1:**
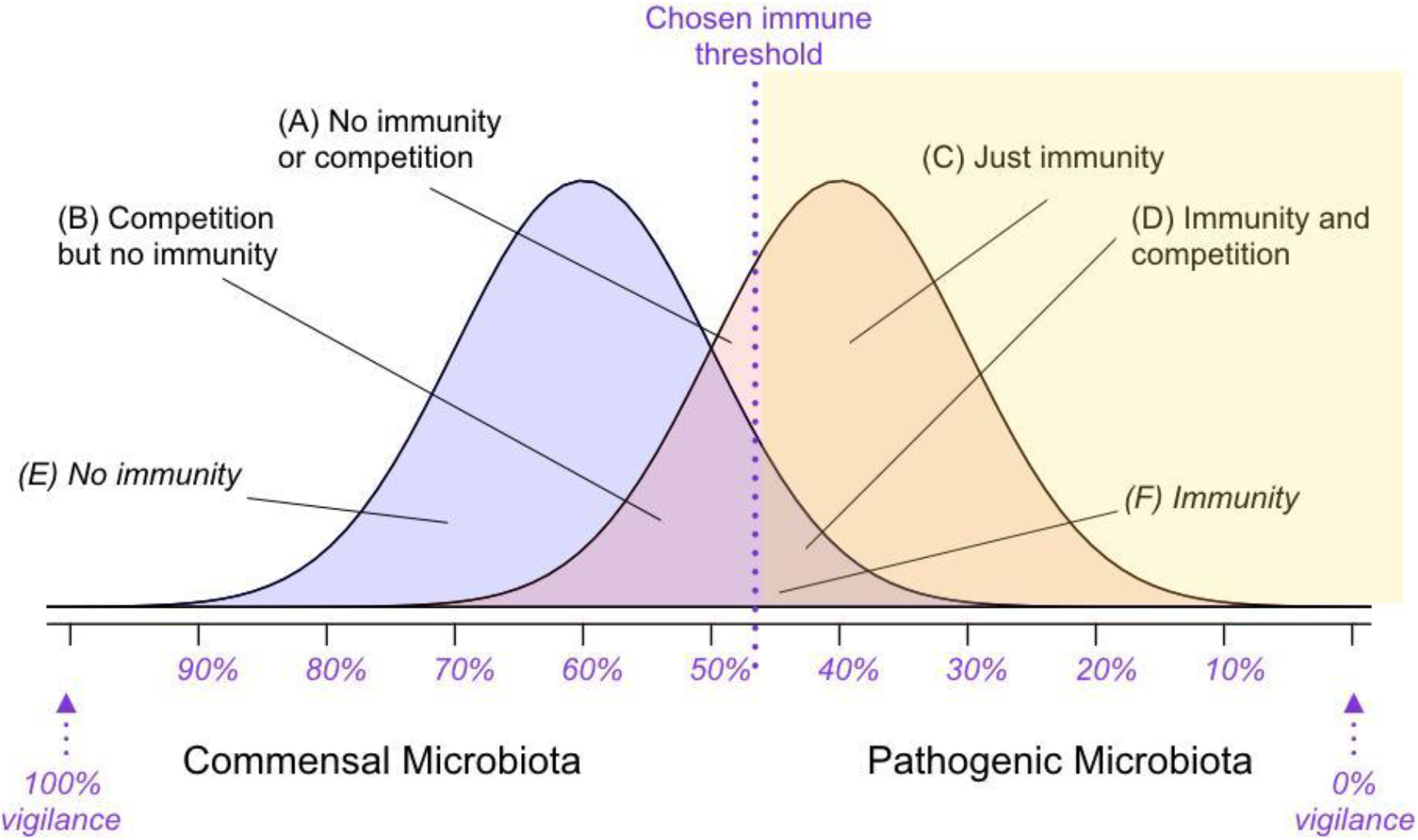
**Schematic of overlap between commensal microbiota (left, blue) and pathogenic microbiota (right, red) communities** along a continuous axis (x axis) reflecting intensity of some immune trigger (e.g., CpG ratio for TLR9). The y axis captures the abundance of individuals corresponding to each point along this continuous axis. The purple line shows the threshold above which an immune response is triggered, and is shaded yellow. Areas of the commensal microbiota distribution above the purple line are ‘false positives’ (inappropriately targeted by the immune system, area defined by *F*); areas of the pathogen distribution below the purple line are ‘false negatives’ (inappropriately ignored by the immune system, areas defined by A and B). Because both the presence or absence of the immune response, but also the presence or absence of the commensal microbiota (via competition) define how pathogenic microbiota affect the host, there are four different categories (A-D, see labels) associated with the pathogen distribution. There are only two associated with the commensal microbiota (E and F), since we are assuming no effect of the pathogen on these species. The optimization problem faced by host immunity is where to draw the immune threshold (purple line) to balance the costs of the pathogenic microbiota and immunity, and the benefits of the commensal microbiota. In this depiction, commensal and pathogenic microbiota communities are shown as having equal abundance, but in reality the pathogenic microbiota might be much rarer, and thus the height of the distribution lower.

This framing reduces the effects of immune discrimination and competition between commensal and pathogenic bacteria to being organized along a single dimension (the x axis on Figure 1); and furthermore assumes that all microbial species classified as ‘commensal’ and all those classified as ‘pathogenic’ have identical effects on host fitness. Both these features are simplifications. If, for example, the beneficial effect of commensal species is highest for those species that most resemble the pathogenic species, this will considerably alter the optimal threshold for an immune response, as it will no longer make sense to sacrifice some of these beneficial species to control the pathogenic species. Nevertheless, it provides a starting point for framing the challenges faced by immunity, and might be appropriate if variation along the scale of discrimination available to the immune system (x axis on Figure 1) overwhelms the individual species differences in terms of effects on the host within the categories commensal and pathogenic microbiota.

### Defining survival probability in the context of a discrimination trade-off

To move from our conceptual mapping of the challenge faced by the immune system (Figure 1) to a measure of host fitness, we express host survival as the outcome of mortality hazards associated with each context. We can express the hazard of mortality for an individual as:

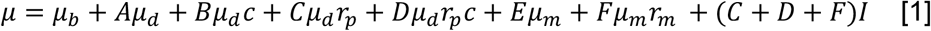

where the quantities *A*, *B*, *C*, *D*, *E*, and *F* refer to areas as shown on Figure 1 (*A*+*B*+*C*+*D* captures the areas occupied by the pathogenic microbiota, and *E*+*F* areas occupied by the commensal microbiota); *μ*_*b*_ is a baseline mortality hazard, *μ*_*d*_ is the mortality hazard associated with the presence of the pathogenic species (and corresponds to their ‘virulence’), *μ*_*m*_ is the mortality hazard associated with the presence of the commensal microbiota (and is either zero or negative).The parameter *r*_*p*_ captures the host’s resistance to the pathogenic microbiota, e.g., its ability to reduce the mortality hazard associated with the presence of pathogenic species to the right of the immune threshold line. This parameter is contained between 0 and 1, where *r*_*p*_ = 1 indicates no effect of the host on the pathogenic species’ impact, and *r*_*p*_ = 0 indicates that the host completely eliminates the effects of the pathogenic species. For completeness, *r*_*m*_captures how much the host reduces the effect of the commensal microbiota. Where the microbiome has beneficial effects on survival (*μ*_*m*_ < 0) this will translate into the host tolerating their commensal microbiota if *r*_*p*_ = 1, and eliminating the beneficial effects if *r*_*p*_ = 0. The parameter *c* captures the degree to which the presence of the commensal microbiota reduces the mortality hazard by the pathogenic microbiota (e.g., by outcompeting or interfering with them) in a manner dependent on their overlap along the continuous axis used by the host. This parameter *c* is also contained between 0 and 1, with larger values indicating less effective competition by the microbiome. The parameter *I* captures the cost of an immune response, and is multiplied by all contexts where an immune reaction has been triggered (e.g., area to the right of the vertical line, shaded yellow, Figure 1).

To evaluate the outcome of different scenarios for the host, each mortality hazard must be multiplied by the relevant area reflecting the combination of the distribution of commensal and pathogenic microbiota, and the positioning of the immune threshold, as labelled on Figure 1; with A, the area below the threshold for immunity consisting of only the pathogen; B the area below the threshold where the pathogenic microbiota overlaps with the commensal microbiota; C, the area above the threshold where only the pathogenic microbiota is present; D, the area above the threshold where both pathogenic and commensal microbiota are present (Figure 1); E, the area where the commensal microbiota is below the threshold; and F, the area where the commensal microbiota is above the threshold and thus potentially eliciting an immune response. An individual’s probability of survival, as a function of the presence of the pathogen and its immune threshold can then be expressed by *s* = *exp*(−*μ*).

### Selection on the immune threshold

Assuming that fertility is not affected by either the microbiota or the immune strategy adopted by the host, maximising survival will maximize host fitness. We can evaluate evolutionary outcomes schematically, mapping out how different contributions to the hazard change as the threshold moves from left (complete immune vigilance) to right (no immune vigilance), and from this, characterize how increases in each of the contributions to the mortality hazard modulates the optimal immune threshold, which is defined by the strategy corresponding to the lowest summed hazard and thus highest survival (Figure 2). This indicates that high costs of immunity, or positive effects of the microbiome could shift the optimal towards ‘no vigilance’ (i.e., drive hosts towards the evolution of no immunity). Indirect effects of the commensal microbiota on pathogen growth, for example, whereby commensal microbiota competition reduces pathogen hazard, could also have this effect.

**Figure 2:**
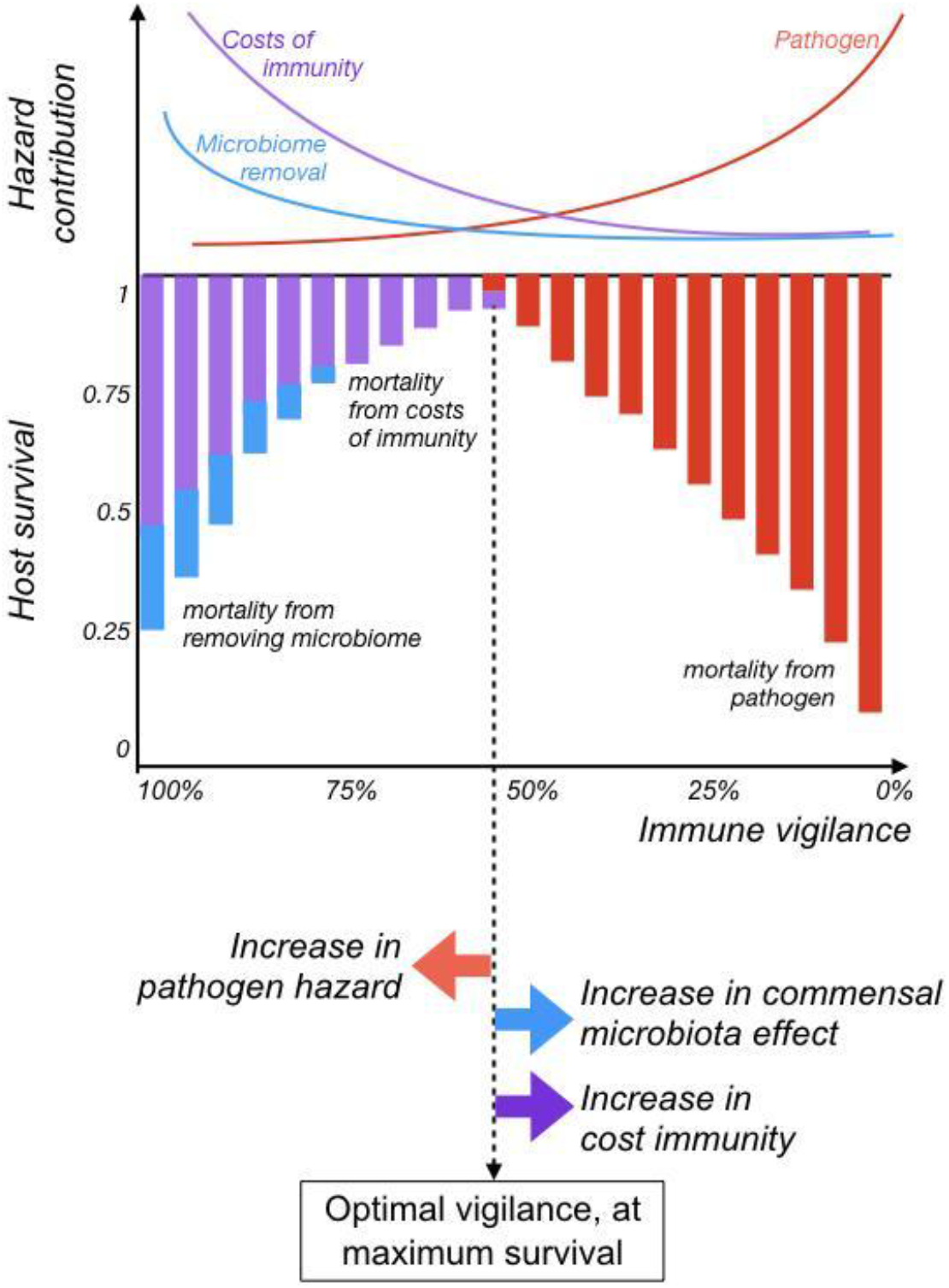
**Schematic relationship between immunity and fitness,** as a function of the position of the immune threshold (x axis), where a threshold on the far left reflects *total vigilance* (points all along the axis trigger a response), and to the far right reflect *no vigilance* (no points along the x axis trigger a response). Contributions to the hazard (y axis, top panel) from pathogenic microbiota (red) increase as immune vigilance declines; immune costs (purple) decrease as immune vigilance declines; and hazards associated with loss of a beneficial microbiota (blue line) decrease as immune vigilance declines. Each of these hazards reduces overall survival additively (y axis, middle panel); and fitness peaks when the sum the various contributions to the hazards is the smallest (dashed vertical line). Increases in pathogenic microbiota related hazards (i.e., vertical red bars falling lower) will shift the immune threshold left (red arrow, bottom panel), increasing the optimal vigilance. Conversely, increases in the costs of immunity, or loss of impacts of a beneficial microbiota will shift the threshold to the right, thus decreasing optimal vigilance (purple and blue arrows, respectively). Competition (parameter *c*) and immune effects (*r*_*p*_ and *r*_*m*_) are not illustrated here because their effect may be context specific (Figure 3).

Using basic probability, we can calculate the areas A-F. For simplicity, we can first evaluate outcomes where the two distributions are perfectly overlapping. This also captures the scenario where the pathogenic microbiota has evolved to be indistinguishable from the commensal microbiota. In this situation, *A* = *C* = 0; *B* = *E* and *D* = *F* = 1 − *B*. The equation becomes:

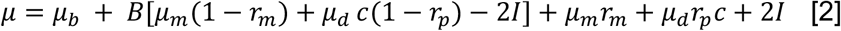

As the immune threshold moves from left to right, reflecting a transition from complete vigilance by the immune system to no vigilance at all, the magnitude of the area denoted *B* increases (Figure 1). As a result, the sign of the factor multiplying *B* will determine whether the outcome of evolution in the presence of a commensal microbiota is effectively the absence of an immune response or a complete immune response (we expect purely binary outcomes in this simple scenario). From this, selection for no vigilance requires that:

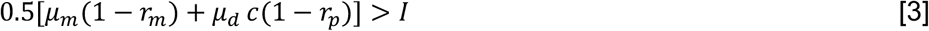

This relationship indicates that the cost of immunity (*I*) required to drive evolution in the host towards a strategy of no immune vigilance will increase: (i) as the hazards increase (whether associated with the commensal microbiota or the pathogenic microbiota); (ii) when the effect of competition from the commensal microbiota on the pathogenic microbiota is reduced (reflected by increased values of *c*); or (iii) if the impact of immunity on the hazards is increased (reflected by decreased values of either *r*_*m*_ or *r*_*p*_).

Focusing on the magnitude of immune cost (*I*) required to drive complete loss of immune vigilance provides a tractable focus for understanding life history constraints (given the array of potential parameters), as well as being of biological interest. Extending results presented in eqn [3] numerically (Figure 3A), we can plot the magnitude of the cost of immunity (*I*) required for evolution to the point of no immune vigilance across a spectrum of commensal and pathogenic microbiota-mediated hazards, and across a range of overlaps between commensal and pathogenic microbiota (Figure 3B, x axis). The required immune cost associated with loss of all immune vigilance increases as overlap declines (Figure 3B, y axis), with higher costs required for lower pathogen hazards (Figure 3B, colours), or in the presence of a commensal microbiota (Figure 3B, line types); and costs decline with diminishingly effective pathogen immunity *r*_*p*_ (Figure 3B, left to right panels).

**Figure 3:**
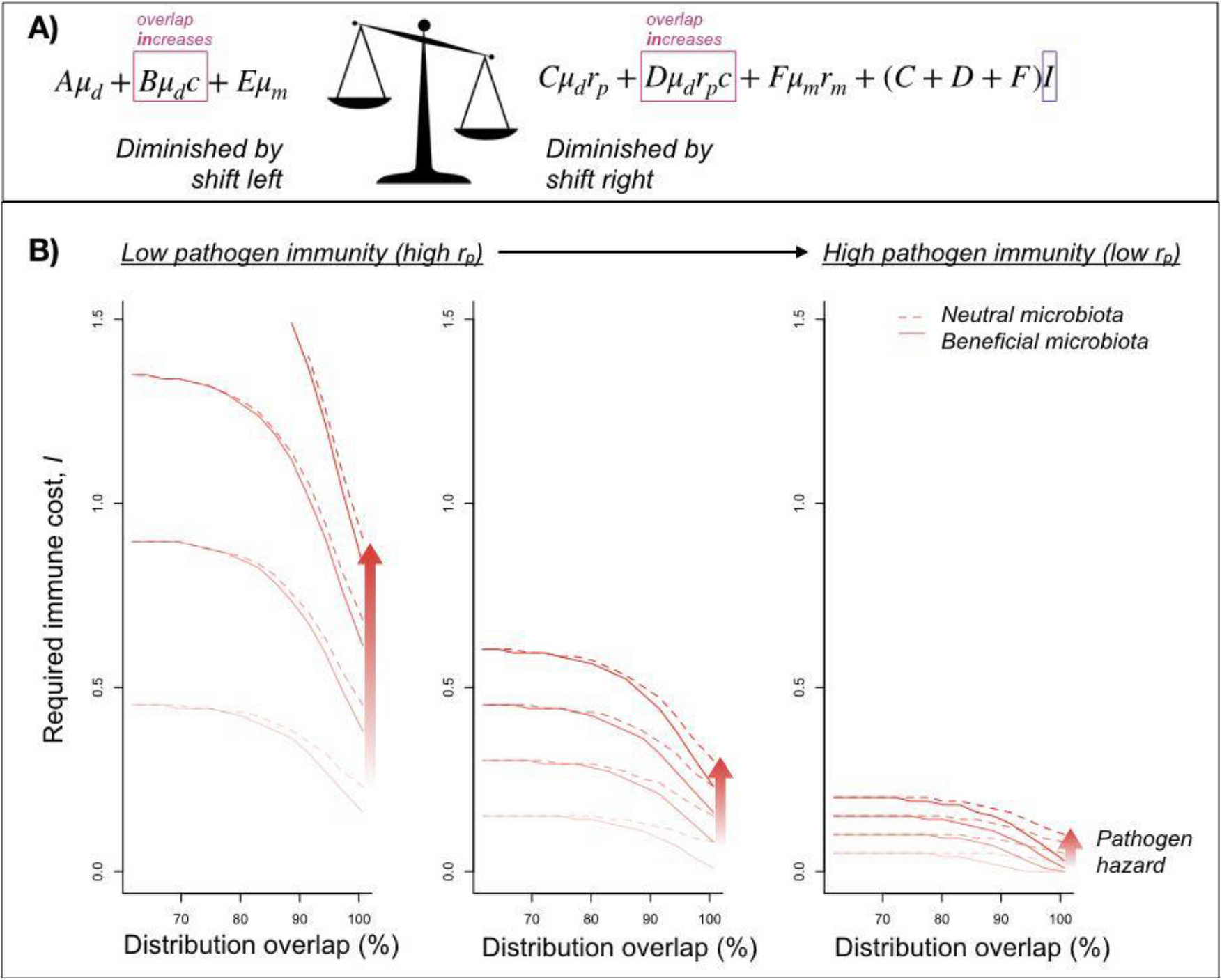
**Cost of immunity required to drive evolution of loss of discrimination** A) This cost is the value of *I* (purple box) required to tip the quantities on the right of the scales (i.e., the components of the hazard that decline as the immune threshold moves right and immune vigilance falls), below the quantities on the left (i.e., components that increase in value as the immune threshold moves right). The effect of overlap between commensal and pathogenic microbiota communities, governed by the competition parameter, *c*, contributes to both sides of the scale (red boxes). B) As the overlap between commensal and pathogenic microbiota communities increases (x axis shows proportion overlap between the two distributions), the cost of immunity required to drive evolution to no immune vigilance (y axis) decreases. All else equal, the cost is greatest for large pathogen hazards (darker red colours) in the absence of commensal microbiota (solid line corresponding to *μ*_*m*_ = −0.2; with *μ*_*m*_ = 0 for the dashed line; other parameters are *r*_*m*_ = 0.3, *c* = 0.5). Reducing the effectiveness of immunity in reducing the hazard associated with pathogen microbiota (by moving from *r*_*p*_ = 0.1 left panel, to *r*_*p*_ = 0.7 right panel, corresponding to ineffectual immunity) results in lower costs allowing loss of host immunity (far right). If the pathogenic microbiota community is rarer than the commensal microbiota community (corresponding to a reduction in the height of the pathogen distribution in Figure 1) this further reduces the cost of immunity required to drive loss of immune vigilance.

How will competition shape the evolution of immune vigilance? To explore this, we can evaluate survival at a range of immune thresholds in the absence (*c* = 1, Figure 4, left) versus presence (*c* = 0, FIgure 4, right) of competition across a spectrum of pathogenic microbiota hazard and immune effectiveness against pathogens (Figure 4, line types, text) for different costs of immunity (purple colours indicate higher costs). If mortality caused by pathogenic microbiota is low, and/or immune effectiveness is high, the optimal strategy is no immune vigilance (Figure 4, dashed line peaks to the right) and is not much modulated by competition (compare left and right panels). For intermediate pathogenic microbiota mortality / immune effectiveness, the optimal strategy is complete vigilance in the absence of competition (Figure 4, left, dotted line peaks to the left). The presence of competition between the commensal and pathogenic microbiota increases host survival across the range of immune vigilance, and slightly reduces the optimal immune vigilance (Figure 4, right, dotted lines increase with the dashed lines then decline at an intermediate threshold). For large pathogen mortality / low immune effectiveness, in the absence of competition, there is little difference in survival across a range of immune thresholds: complete vigilance results in very minor increases in survival for low costs of immunity, and vice versa (Figure 4, left solid line). The presence of competition results in larger survival across the board, and further, alters the optimal to being no immune vigilance, with this reflecting a considerable survival advantage (Figure 4, right, solid line peaks to the right).

**Figure 4:**
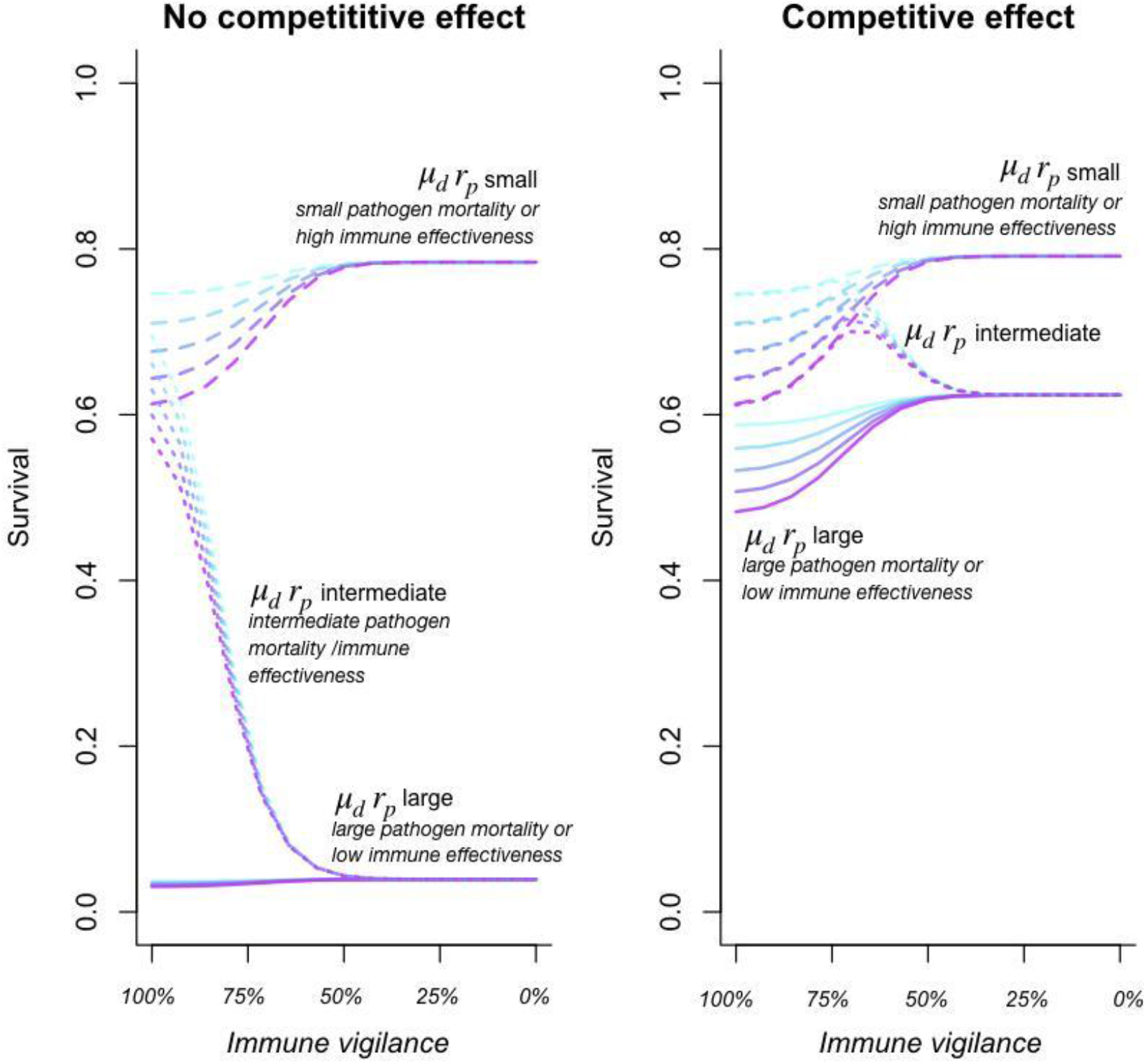
**Effect of competition on survival** (y axis) across a range of levels of immune vigilance (x axis) in the absence (left, *c* = 1) and presence (right, *c* = 0) of competition for different combinations (text, line types) of the product of pathogen mortality (*μ*_*d*_ = 0.01 or *μ*_*d*_ = 3) and immune effectiveness (*r*_*p*_ = 0.01 or *r*_*p*_ = 1) for different costs of immunity (*I* = 0.1 shown in light blue is the lowest cost; through to *I* = 0.2, in purple as the highest cost, i.e., corresponding to the greatest reduction in survival). The presence of competition increases survival (right hand plot vs. left), but also can qualitatively change the outcome of selection on immune vigilance. For large pathogen microbiota associated mortality and/or low immune effectiveness (solid line) the outcome switches from close to no directional selection on vigilance to strong selection for no vigilance (solid lines, compare left and right), while for intermediate pathogen mortality and/or immune effectiveness, selection shifts from favouring complete vigilance to intermediate vigilance (dotted line, compare left and right).

In results presented to this point, we have effectively assumed that the abundance of commensal and pathogenic microbiota are equal. In reality, pathogenic species are likely to be considerably more rare. Altering results to encompass lower abundance in the pathogenic microbiota community (equivalent to reducing the height of the red distribution on Figure 1) results in a qualitatively similar results, but with loss of immune vigilance achieved at lower levels for comparable parameter sets.

### Dysbiosis at high or low diversity

In many settings, reduced microbiome diversity has been linked to ‘dysbiosis’, or reduced health as a result of perturbed microbiome ecology. However, in others, such as the vaginal microbiome, evidence suggests that reduced diversity is in fact a sign of health. To better capture how the impacts of microbiome diversity shape selection on immune system response, we assume that the range of diversity left intact in the wake of immune activity (e.g., values to the left of the threshold for the immune response, Figure 1) affects the overall hazard. Diversity is measured using Simpson’s diversity index,

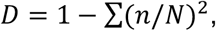

where *n* is the number of individuals of each species (identified as positions along the x axis on Figure 1) and *N* is the total number of individuals, in both cases taking only points below the threshold. This index thus represents the probability that two individuals randomly selected from a sample will belong to different species, and is thus essentially a saturating function of the position of the threshold from left to right, saturating at 1.

The effect of diversity on the hazard *μ*_*m.div*_ may be either negative or positive, and thus may increase or decrease the mortality hazard. The expression for the mortality hazard becomes:

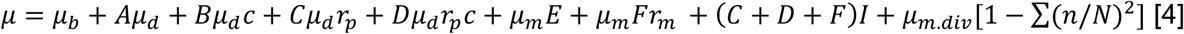

Conceptually, this corresponds to adding one more curve to the schematic of hazards in Figure 2, either increasing (where the hazard increases with diversity) or decreasing (where the hazard decreases with diversity). This illustrates that where dysbiosis increases as diversity declines (e.g., as in (Lawley et al. 2012)), this will select for a decline in immune vigilance; and vice versa.

Again, as discussed in introducing the framework (Figure 1), this neglects the important fact that some species may be far more important than others in terms of their effects on the host; and as in the previous case, how diversity shapes the optimal immune discrimination might be substantially altered by this.

## Discussion

We provide a first framing of how the presence of microbiome species might modulate the evolution of innate immunity where the immune system faces the challenge of discriminating between pathogenic and other microbe species (Figure 1). We separate out direct positive effects of the microbiome on reducing mortality (even in the absence of infection) with indirect effects from competing with and reducing mortality effects of pathogenic species (captured by the parameter *c*). We show that either effect can shift the evolution of host immunity towards loss of vigilance; that the effects are amplified as similarity between the microbiome and pathogen communities increases (Figure 3B, x axis); and that when immune effectiveness against the pathogen is weak (Figure 3B, right panel) immunity can be lost even when not very costly. The likelihood of loss of immune vigilance will be further amplified if dysbiosis declines with diversity.

Our framework makes explicit some of the core trade-offs which define whether microbiome species presence can drive loss of immune vigilance. A comparative framing provides one possible approach to testing the resulting predictions. Evidence for protective effects of the microbiome exists across a range of hosts, including against *Pseudomonas syringae* in horse chestnut trees (Koskella et al. 2017), tomato (Berg and Koskella 2018) and arabidopsis plants (Innerebner, Knief, and Vorholt 2011); against both salmonella (Kubinak et al. 2015) and intestinal infection with *Entamoeba histolytica (Watanabe et al. 2017)* in mice; and against the conjunctival bacterial pathogen *Mycoplasma gallisepticum* in house finches (Thomason et al. 2017). Furthermore, the role of individual defensive symbionts on host fitness has been demonstrated both empirically (Vorburger and Perlman 2018) and theoretically (King and Bonsall 2017). Our framework predicts that, all else equal, the relevant host species (or genotype) will have diminished immune discrimination against pathogens that most resemble protective bacteria, for example reduced responsiveness of pattern recognition receptors (Jones and Dangl 2006; Trdá et al. 2014).

Although we discuss our results in microbiome community terms, where all commensal and pathogenic microbiota have the same (positive or negative) effect on the host, our broad conclusions will hold for a single pathogenic species, with some small distribution along the possible immune trigger, or a single commensal species, likewise. The tension between paying the costs of immunity and those of harbouring pathogenic microbiota will play out even if the non-pathogenic microbiota considered are neutral (Figure 3). For tractability, our framing also collapses the possible triggers for immunity to a single axis (Figure 1). The molecular patterns recognized by immunity’s Pattern Recognition Receptors are by definition constrained, so varied receptor responsiveness (Hawn et al. 2007; Cecil et al. 2016; Trdá et al. 2014; Vetter et al. 2012) might be low dimensional. Functional characteristics indicative of pathogen’s presence (e.g., deviations from homeostasis) are likely to be similar across a broad array of pathogens (DeYoung and Innes 2006; Chovatiya and Medzhitov 2014), also suggesting low dimensionality. Both these aspects suggest that although a single dimension may not perfectly reflect how microbial species trigger immunity, it provides a reasonable starting point; albeit one perhaps most relevant considering interactions happening within kingdoms (bacterial pathogens with bacterial microbiome, viral pathogens with virome, and fungal pathogens with mycobiome), given higher potential for overlap of shared immune mechanisms).

Our primary focus here is on innate immunity, but the front line of defense in many organisms, for many species, is adaptive immunity, especially in protecting hosts from re-exposure to parasites. Nevertheless, cross-talk with innate immunity remains a crucial element in the activation of adaptive immunity (Iwasaki and Medzhitov 2015), so the trade-offs introduced here are likely to still be relevant. Furthermore, this framing raises interesting questions about how the challenge of distinguishing pathogenic from neutral or commensal microbes intersects with the intricate evolution of tolerance of microbiome species displayed by adaptive immunity in vertebrates (Cebula et al. 2013). On the time-scales of host ontogeny, introduction of commensal species outside of a critical window of exposure could lead to a host either excluding a potentially protective microbiome species, or mounting an immune defense against non-pathogenic bacteria in a way that leads to autoimmunity (Gensollen et al. 2016).

Across the field, a focus on host evolution relative to pathogen (or microbiome) characteristics remains relatively rare, as the shorter time-scales of microbe generations suggest much more rapid selection. Nevertheless, hosts indubitably do respond to pathogen selection, and by focusing on highly conserved aspects of microbe molecular make-up, we narrow the time-scale mismatch in this co-evolutionary process. Yet, while our focus is on host evolution, we can also evaluate our results in the context of microbe evolution. In particular, the situation of completely overlapping pathogen and microbiome species distributions (equation [3]) captures the situation where the pathogen has evolved to be completely indistinguishable from the microbiome. Another potential direction for coevolution is selection for signaling by commensal microbiota to facilitate recognition by adaptive immunity (Ost and Round 2018), equivalent to increasing the separation between the two distributions in Figure 1. Both microbiome and pathogen species might also be able to evolve to evade immune detection altogether. Modifying our model to include a lower bound on where the immune threshold can be drawn (e.g., in Figure 1, the vertical purple line can no longer go all the way to the right) will account for this by bounding the optimal strategy that the host can evolve - qualitative predictions of our model are unchanged.

Longevity or ‘pace of life’ is often a focal explanatory variable when evaluating the evolution of immunity (Martin, Weil, and Nelson 2007). Here, if the abundance of commensal and pathogenic microbiota is constant across the life-cycle, then we expect no changes in strategy with longevity, since all extra hazards are essentially an extra extrinsic mortality hazard (Caswell 2007). However, in reality, rare microbiota may be less likely to colonize hosts, resulting in a later age of their acquisition. Thus, if pathogenic microbiota are rare, they are likely to be acquired after commensal or neutral microbiota. Given the potential for competition from already-present commensal or neutral microbiota, and the ubiquitous effect of declining selection pressures with age (Hamilton 1966), this could amplify selection for loss of immune vigilance. Yet, longevity is likely to intersect with many other aspects of microbe ecology. In particular, longer lived species might encounter a higher diversity of microbes, simply given longer exposure times during which microbe colonization can occur. Where most microbial species have neutral effects, longer lived species are predicted to retain immune vigilance, all else being equal, since the cumulative costs of pathogens (even if rare) will translate into a significant reduction in long-lived host reproductive value relative to shorter lived species. An increase of dysbiosis with microbe diversity will tend to amplify this effect (noting, however, that the reverse pattern is often reported). Since many features, including costs of immunity might also covary with longevity, careful titration of selection pressures will be required to evaluate the strength of this prediction in natural systems.

Since protective effects of microbiome species are predicted to drive the loss of host immune vigilance, hosts could be left vulnerable to attack by novel pathogens that too closely resemble allies that the host has been selected to ignore. Conversely, if the presence of protective bacteria actually covaries with the presence of pathogens, it has recently been shown that this can result in increased host investment in resistance occurring in tandem with the presence of protective symbionts (Hrček et al. 2018), again emphasizing that the details of microbial ecology matter.

To conclude, while our framework makes a number of simplifying assumptions (e.g., ignoring immune responses triggered by tissue-specific damage, a widely observed phenomenon; and collapsing of potentially multidimensional immune discrimination), it makes a first set of predictions about optimal immune discrimination, both as a function of overlap between pathogen and microbiome, and of microbiome-mediated protection. This initial framing lays a foundation for future work exploring the under-studied question of selection on host immunity in light of our expanding understanding of the microbiome. It also brings to light the need for more explicit data examining the overlap in host recognition of pathogens and commensal microbiota.

## Author Contributions

CJEM and BK developed the concept; CJEM developed the models; CJEM and BK wrote the paper.

## Notes

#### Summary of Updates

Clarity, detail, and improved figures

## References

Berg, Maureen, and Britt Koskella. (2018). “Nutrient- and Dose-Dependent Microbiome-Mediated Protection against a Plant Pathogen.” Current Biology: CB 28 (15): 2487–92.e3.

Caswell, Hal. (2007). “Extrinsic Mortality and the Evolution of Senescence.” Trends in Ecology & Evolution 22 (4): 173–74.

Cebula, Anna, Michal Seweryn, Grzegorz A. Rempala, Simarjot Singh Pabla, Richard A. McIndoe, Timothy L. Denning, Lynn Bry, Piotr Kraj, Pawel Kisielow, and Leszek Ignatowicz. (2013). “Thymus-Derived Regulatory T Cells Contribute to Tolerance to Commensal Microbiota.” Nature 497 (7448): 258–62.

Cecil, Jessica D., Neil M. O’Brien-Simpson, Jason C. Lenzo, James A. Holden, Yu-Yen Chen, William Singleton, Katelyn T. Gause, Yan Yan, Frank Caruso, and Eric C. Reynolds. (2016). “Differential Responses of Pattern Recognition Receptors to Outer Membrane Vesicles of Three Periodontal Pathogens.” PloS One 11 (4): e0151967.

Chovatiya, Raj, and Ruslan Medzhitov. (2014). “Stress, Inflammation, and Defense of Homeostasis.” Molecular Cell. 54(2): 281–288.

DeYoung, Brody J., and Roger W. Innes. (2006). “Plant NBS-LRR Proteins in Pathogen Sensing and Host Defense.” Nature Immunology 7 (12): 1243–49.

Felix, Georg, Juliana D. Duran, Sigrid Volko, and Thomas Boller. (1999). “Plants Have a Sensitive Perception System for the Most Conserved Domain of Bacterial Flagellin.” The Plant Journal: For Cell and Molecular Biology 18 (3): 265–76.

Friesen, Maren L., Stephanie S. Porter, Scott C. Stark, Eric J. von Wettberg, Joel L. Sachs, and Esperanza Martinez-Romero. (2011). “Microbially Mediated Plant Functional Traits.” Annual Review of Ecology, Evolution, and Systematics 42 (1): 23–46.

Gensollen, Thomas, Shankar S. Iyer, Dennis L. Kasper, and Richard S. Blumberg. (2016). “How Colonization by Microbiota in Early Life Shapes the Immune System.” Science 352 (6285): 539–44.

Gómez-Gómez, Lourdes, and Thomas Boller. (2002). “Flagellin Perception: A Paradigm for Innate Immunity.” Trends in Plant Science 7 (6): 251–56.

Hamilton, W. D. (1966). “The Moulding of Senescence by Natural Selection.” Journal of Theoretical Biology. https://doi.org/10.1016/0022-5193(66)90184-6.

Hanssen, S. A., D. Hasselquist, I. Folstad, and K. E. Erikstad. (2004). “Costs of Immunity: Immune Responsiveness Reduces Survival in a Vertebrate.” Proceedings of the Royal Society B: Biological Sciences 271 (1542): 925–30.

Hawn, Thomas R., E. Ann Misch, Sarah J. Dunstan, Guy E. Thwaites, Nguyen T. N. Lan, Hoang T. Quy, Tran T. H. Chau, et al. (2007). “A Common Human TLR1 Polymorphism Regulates the Innate Immune Response to Lipopeptides.” European Journal of Immunology 37 (8): 2280–89.

Hrček, Jan, Benjamin J. Parker, Ailsa H. C. McLean, Jean-Christophe Simon, Ciara M. Mann, and H. Charles J. Godfray. (2018). “Hosts Do Not Simply Outsource Pathogen Resistance to Protective Symbionts.” Evolution. 72(7): 1488–1499.

Innerebner, Gerd, Claudia Knief, and Julia A. Vorholt. (2011). “Protection of Arabidopsis Thaliana against Leaf-Pathogenic Pseudomonas Syringae by Sphingomonas Strains in a Controlled Model System.” Applied and Environmental Microbiology 77 (10): 3202–10.

Iwasaki, Akiko, and Ruslan Medzhitov. (2015). “Control of Adaptive Immunity by the Innate Immune System.” Nature Immunology 16(4): 34.

Jaenike, John. (2012). “Population Genetics of Beneficial Heritable Symbionts.” Trends in Ecology & Evolution 27 (4): 226–32.

Jones, Jonathan D. G., and Jeffery L. Dangl. (2006). “The Plant Immune System.” Nature 444(7117): 323.

King, Kayla C., and Michael B. Bonsall. (2017). “The Evolutionary and Coevolutionary Consequences of Defensive Microbes for Host-Parasite Interactions.” BMC Evolutionary Biology 17 (1): 190.

Koskella, Britt, Lindsay J. Hall, and C. Jessica E. Metcalf. (2017). “The Microbiome beyond the Horizon of Ecological and Evolutionary Theory.” Nature Ecology & Evolution 1 (11): 1606–15.

Koskella, Britt, Sean Meaden, William J. Crowther, Roosa Leimu, and C. Jessica E. Metcalf. (2017). “A Signature of Tree Health? Shifts in the Microbiome and the Ecological Drivers of Horse Chestnut Bleeding Canker Disease.” The New Phytologist 215 (2): 737–46.

Kubinak, Jason L., W. Zac Stephens, Ray Soto, Charisse Petersen, Tyson Chiaro, Lasha Gogokhia, Rickesha Bell et al. (2015). “MHC variation sculpts individualized microbial communities that control susceptibility to enteric infection.” Nature communications 6: 8642.

Lawley, Trevor D., Simon Clare, Alan W. Walker, Mark D. Stares, Thomas R. Connor, Claire Raisen, David Goulding, et al. (2012). “Targeted Restoration of the Intestinal Microbiota with a Simple, Defined Bacteriotherapy Resolves Relapsing Clostridium Difficile Disease in Mice.” PLoS Pathogens 8 (10): e1002995.

Leung, Chung-Yin, and Joshua S. Weitz. (2019). “Not by (Good) Microbes Alone: Towards Immunocommensal Therapies.” Trends in Microbiology 27 (4): 294–302.

Levy, Asaf, Isai Salas Gonzalez, Maximilian Mittelviefhaus, Scott Clingenpeel, Sur Herrera Paredes, Jiamin Miao, Kunru Wang, et al. (2018). “Genomic Features of Bacterial Adaptation to Plants.” Nature Genetics 50 (1): 138–50.

Littman, Dan R., and Eric G. Pamer. (2011). “Role of the Commensal Microbiota in Normal and Pathogenic Host Immune Responses.” Cell Host & Microbe 10 (4): 311–23.

Martin, Lynn B., 2nd, Zachary M. Weil, and Randy J. Nelson. (2007). “Immune Defense and Reproductive Pace of Life in Peromyscus Mice.” Ecology 88 (10): 2516–28.

Ost, Kyla S., and June L. Round. (2018). “Communication Between the Microbiota and Mammalian Immunity.” Annual Review of Microbiology 72 (September): 399–422.

Pohar, Jelka, Duško Lainšček, Ryutaro Fukui, Chikako Yamamoto, Kensuke Miyake, Roman Jerala, and Mojca Benčina. (2015). “Species-Specific Minimal Sequence Motif for Oligodeoxyribonucleotides Activating Mouse TLR9.” Journal of Immunology 195 (9): 4396–4405.

Sheldon, B. C., and S. Verhulst. (1996). “Ecological Immunology: Costly Parasite Defences and Trade-Offs in Evolutionary Ecology.” Trends in Ecology & Evolution 11 (8): 317–21.

Snelders, Nick C., Graeme J. Kettles, Jason J. Rudd, and Bart P. H. J. Thomma. (2018). “Plant Pathogen Effector Proteins as Manipulators of Host Microbiomes?” Molecular Plant Pathology 19 (2): 257–59.

Thaiss, Christoph A., Niv Zmora, Maayan Levy, and Eran Elinav. (2016). “The Microbiome and Innate Immunity.” Nature. 535(7610): 65.

Thomason, Courtney A., Nathan Mullen, Lisa K. Belden, Meghan May, and Dana M. Hawley. (2017). “Resident Microbiome Disruption with Antibiotics Enhances Virulence of a Colonizing Pathogen.” Scientific Reports 7 (1): 16177.

Trdá, Lucie, Olivier Fernandez, Freddy Boutrot, Marie-Claire Héloir, Jani Kelloniemi, Xavier Daire, Marielle Adrian, et al. (2014). “The Grapevine Flagellin Receptor VvFLS2 Differentially Recognizes Flagellin-Derived Epitopes from the Endophytic Growth-Promoting Bacterium Burkholderia Phytofirmans and Plant Pathogenic Bacteria.” The New Phytologist 201 (4): 1371–84.

Vetter, M. Madlen, Ilkka Kronholm, Fei He, Heidrun Häweker, Matthieu Reymond, Joy Bergelson, Silke Robatzek, and Juliette de Meaux. (2012). “Flagellin Perception Varies Quantitatively in Arabidopsis Thaliana and Its Relatives.” Molecular Biology and Evolution 29 (6): 1655–67.

Vogel, Christine, Natacha Bodenhausen, Wilhelm Gruissem, and Julia A. Vorholt. (2016). “The Arabidopsis Leaf Transcriptome Reveals Distinct but Also Overlapping Responses to Colonization by Phyllosphere Commensals and Pathogen Infection with Impact on Plant Health.” The New Phytologist 212 (1): 192–207.

Vorburger, Christoph, and Steve J. Perlman. (2018). “The role of defensive symbionts in host– parasite coevolution.” Biological Reviews 93(4): 1747–1764.

Watanabe, Koji, Carol A. Gilchrist, Md Jashim Uddin, Stacey L. Burgess, Mayuresh M. Abhyankar, Shannon N. Moonah, Zannatun Noor, et al. (2017). “Microbiome-Mediated Neutrophil Recruitment via CXCR2 and Protection from Amebic Colitis.” PLoS Pathogens 13 (8): e1006513.

